# Intestinal lamina propria supports acquired eTreg suppressor function

**DOI:** 10.1101/2022.08.26.505428

**Authors:** Yisu Gu, Raquel Bartolomé-Casado, Chuan Xu, Alina Janney, Cornelia Heuberger, Claire Pearson, Sarah Teichmann, Emily E Thornton, Fiona Powrie

**Affiliations:** Kennedy Institute of Rheumatology, NDORMS, University of Oxford; Wellcome Sanger Institute, Wellcome Genome Campus, Hinxton, Cambridge; Department of Pathology, Oslo University Hospital – Rikshospitalet, Oslo, Norway; Theory of Condensed Matter, Cavendish Laboratory, Department of Physics, University of Cambridge, Cambridge; MRC Human Immunology Unit, MRC Weatherall Institute of Molecular Medicine, University of Oxford; Nuffield Department of Medicine, University of Oxford

**Author notes:** these authors contributed equally (Yisu Gu, Raquel Bartolomé-Casado, Emily E Thornton, Fiona Powrie).

## Abstract

The intestinal immune system must maintain tolerance to commensal microbiota and self antigens whilst defending against invading pathogens. Recognising how homeostasis is established and maintained in a complex immune environment such as the gut is critical to understanding how to re-establish tolerance once broken in inflammatory disorders. Peripherally induced regulatory T cells (Tregs) play a key role in homeostasis. In intestinal tissue, Tregs work in concert with a diverse network of cells but which cellular interactions occur to instruct Treg adaptation and acquisition of distinct Treg suppressor function is not clear. We used two-photon *in vivo* live imaging and NICHE-seq [1] to deep phenotype *Helicobacter hepaticus* (*Hh*)-specific Tregs with shared specificity but distinct spatially compartmentalised functions in the tissue. We show transcriptionally distinct central Treg (cTreg) and effector Treg (eTreg) populations in lymphoid versus gut tissue. The lamina propria (LP), and not embedded lymphoid aggregates (LA), is the key location of acquired immune suppressor eTreg function. Tregs recruited to the LP compartment are the dominant interacting cell type and acquired a more effector Treg profile with upregulation of *Areg, Gzmb, Icos, Tigit, Tnfrsf4* (OX40), and *Tnfrsf18* (GITR). We identify IL-1β^+^ macrophages, CD206+ macrophages, and ILC2 in the LP niche as the key players governing Treg survival and function. In contrast, LA, dominated by interactions with ILC3s and populations of IL-6^+^ DCs, are equipped to tip the balance towards a pro-inflammatory response. By functionally isolating the gut tissue from secondary lymphoid organs, we show that eTregs maintain their phenotype in the context of inflammatory insult. Blocking their key effector molecule, IL-10, results in locally differentiated Th17 cell proliferation without overt inflammation due to local IL-10 independent mechanisms that constrain inflammation. Our results reveal a previously unrecognised spatial mechanism of tolerance, and demonstrate how knowledge of local interactions can guide cell function and potentially be manipulated for the next generation of tolerance-inducing therapies.

## Results

The pathobiont *Helicobacter hepaticus* (*Hh*) establishes life-long infection in the cecum of normal mice. A key host adaptation ensuring immune homeostasis in the face of chronic infection is the production of IL-10 by Tregs [2–4]. Maladaptation of this response results in colitis in mice, and similar processes are thought to underlie very early onset inflammatory bowel disease (VEO-IBD) with deficiencies in the IL-10 pathway a major cause [5]. The MLN has been shown to be a key site of Treg induction [6], but the key anatomical location for induction and maintenance of effector Treg suppressor function is yet to be elucidated. We used HH7-2tg T cells [7] (TCR^Hh^) in *Hh* colonised hosts to follow the natural history of antigen-specific T cells experiencing key interactions and gaining and sustaining Treg effector functions in the tissue microenvironment.

To map where adaptive responses to *Hh* occur, we characterised the intestinal lymphoid and non-lymphoid tissue compartments of WT C57BL/6 mice. Lymphoid tissue comprised, secondary lymphoid organs (SLO) including mesenteric lymph nodes (MLN), the caecal patch (CP) and distal colon organised lymphoid structures (OLS) (Fig 1a). Small lymphoid aggregates (LA) are present in the caecum and proximal colon. LA do not have organised T and B-cell zones, and contain a spectrum of tissue organisation from cell aggregates to cryptopatches and isolated lymphoid follicles (ILF) (Supp Fig 1a-c). Previous tracking of naïve TCR^Hh^ *in vivo* showed differentiation to Rorgt^+^ Foxp3^+^ Tregs in colons, but that work did not exclude OLS from the analysis [7]. We sought to understand the natural history of TCR^Hh^ to *Hh* in the context of lymphoid and non-lymphoid compartments, referring to the ‘tissue’ as the composite of the lamina propria (LP) and the lymphoid aggregates (LA), but exclusive of OLS.

**Figure 1.**
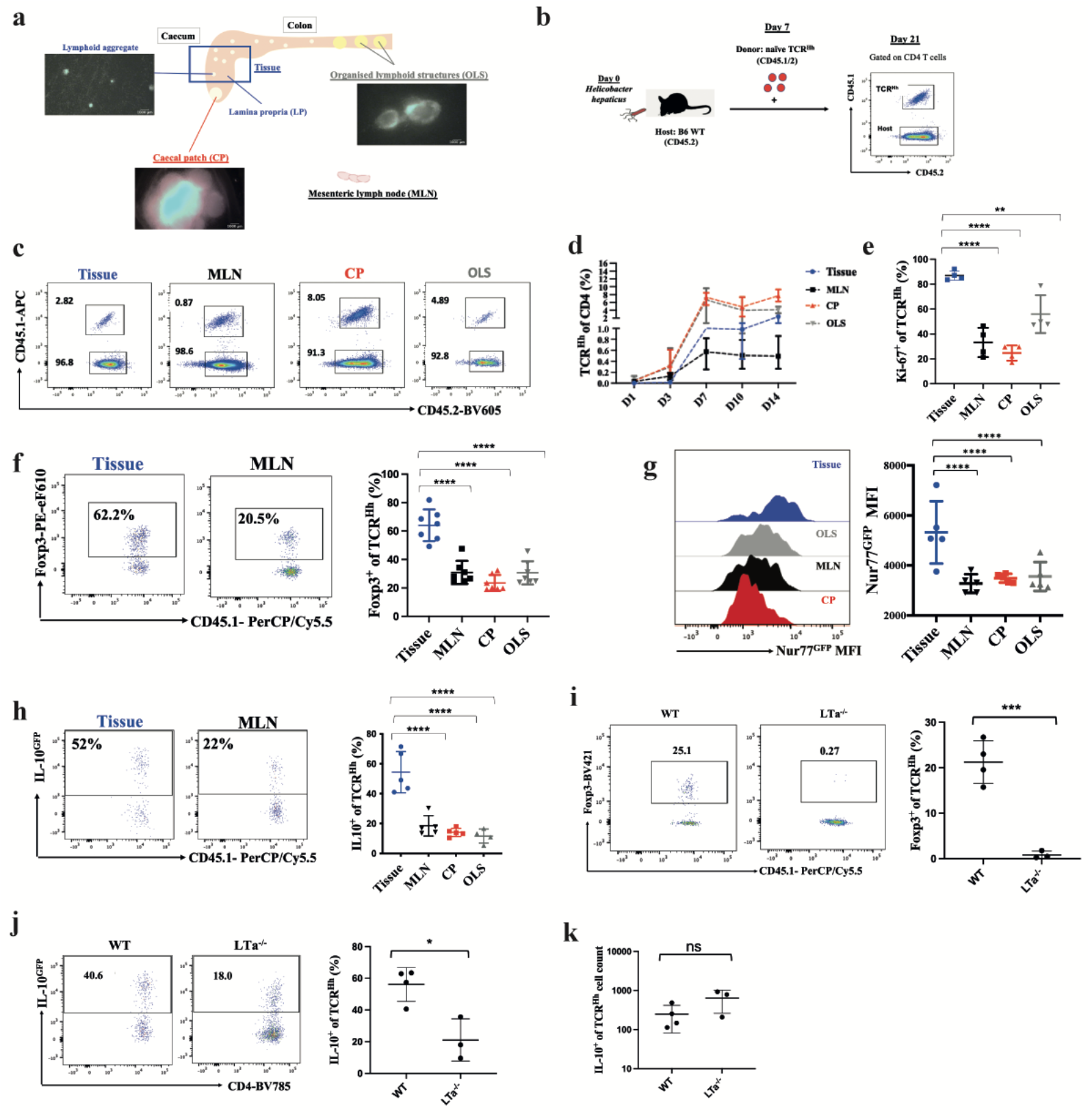
Colon tissue supports the highest IL10 production and strongest TCR signalling in response to *Helicobacter hepaticus*. a. Diagram of tissue and lymphoid structures in the murine gut, with representative images of caecal patch (CP), organised lymphoid structures (OLS) and lymphoid aggregates (LA) in hCD2-DsRed mouse (scale bar 1000μm).
b. Transfer of naïve CD45.1^+^ TCR^Hh^ T cells into *Hh*-colonised hosts.
c. Representative FACS of T TCR^Hh^ 14 days post transfer at the indicated sites.
d. Frequency of TCR^Hh^ among total CD4 T cells at indicated time points post transfer.
e. Frequency of Ki-67^+^ of TCR^Hh^ at 14 days post transfer at the indicated sites.
f. Representative FACS and frequency of Tregs of TCR^Hh^ at 14 days post transfer.
g. Representative histogram overlay and GFP MFI of TCR^Hh^Nur77^GFP^ 11 days post transfer.
h. Representative FACS and frequency of IL-10^+^ among TCR^Hh^IL-10^GFP^ 11 days post transfer.
i. Representative FACS and frequency of Tregs among TCR^Hh^ in tissue of WT and LTα^-/-^ hosts at 10 days post transfer.
j. Representative FACS and frequency of IL-10^+^ among TCR^Hh^ in tissue of WT and LTα^-/-^ hosts at 10 days post transfer.
k. Absolute cell count of IL-10^+^ among TCR^Hh^ in tissue of WT and LTα^-/-^ hosts at 10 days post transfer.

Naive TCR^Hh^ transferred into *Hh*-colonised hosts (Fig 1b and Supp Fig 1d) home to the SLOs within 24 hours and onwards to the tissue approximately 1 week later (Fig 1c-d). Co-transfer of congenically marked TCR^Hh^ and TCR^Hh^ *Rag1*^-/-^ cells confirmed no effect of endogenous TCRa rearrangement (Supp Fig 2a-d). Further, TCR^Hh^ were not detected in un-colonised hosts, demonstrating their dependence on cognate antigen for survival (Supp Fig 2e). To pinpoint where important cellular interactions and activation steps take place, we quantified proliferation, Treg differentiation, TCR signalling and IL-10 production across the tissue compartments. Despite early activation in lymphoid tissues (Supp Fig 2f), at 14 days after transfer, tissue TCR^Hh^ cells are more proliferative than their lymphoid counterparts (Fig 1e), and a larger proportion of TCR^Hh^ cells differentiated into Foxp3+ Tregs (Fig 1f) within tissue than SLOs.

As Treg suppressor function has been shown to depend on the T cell receptor (TCR) [8], we wondered whether ongoing TCR stimulation also occurs in TCR^Hh^ in any of the defined tissue microenvironments. Transfer of naïve T cells from TCR^Hh^ crossed to Nur77^GFP^ mice [10] (TCR^Hh^ Nur77^GFP^) into *Hh*-colonised hosts showed the proportion expressing GFP and GFP MFI as a measure of TCR signalling to be highest in the tissue at 11 days post transfer (Fig 1g). Transfer of TCR^Hh^ crossed to B6.Foxp3^hCD2^ IL-10^GFP^ reporter mice [9] (TCR^Hh^IL-10^GFP^) also revealed the tissue as the site of the highest proportion of IL-10^+^ TCR^Hh^ cells (Fig 1h). To determine whether TCR engagement and IL-10 production were due to local stimulation, we used the sphingosine 1-phosphate receptor agonist FTY720 to block lymph node egress following TCR^Hh^ recruitment into the gut to ensure detection of local GFP induction (Supp Fig 3a-c). Again, the strongest TCR signalling and highest IL-10 production was detected in tissue TCR^Hh^ cells, which were on par with controls (Supp Fig 3d-g). Absolute numbers of tissue TCR^Hh^ were comparable between FTY720 treated and controls, suggesting local proliferation of TCR^Hh^ (Supp Fig 3h). These data point to the gut tissue as the dominant site of ongoing Treg activation and effector function in the maintenance of tolerance to the pathobiont *Hh*. In fact, lymphotoxin-a deficient mice (L.Tα^-/-^) that are devoid of MLNs, CP, OLS and LA (Supp Fig 4a-b, and data not shown) [10] still supported TCR^Hh^ homing into the tissue (Supp Fig 4c-e). However, in the absence of lymph nodes, TCR^Hh^ did not differentiate into Tregs (Fig 1i), indicating a non-redundant role for SLOs in Treg differentiation. TCR^Hh^ cells in LTα^-/-^ hosts retained the ability to make IL-10 in similar absolute numbers, albeit at a lower frequency compared to controls (Fig 1j-k, Supp Fig 4f). These data show SLOs are critical for Treg induction, but interactions in the tissue can induce non-Treg TCR^Hh^IL-10 production independent of SLOs and prevent colitis (Supp Fig 4g).

Our results demonstrate tissue interactions play a dominant and previously unrecognised role in TCR stimulation and IL-10 production of microbe-reactive T cells, including Tregs. We next wished to determine if the LP and LA niches played individual roles in shaping the TCR^Hh^ response to *Hh*. We bred TCR^Hh^Nur77^GFP^ and TCR^Hh^IL-10^GFP^ to hCD2-DsRed mice [11] (TCR^Hh^CD2^Dsred^Nur77^GFP^ or TCR^Hh^CD2^Dsred^IL10^GFP^) and separately transferred naïve T cells from these reporter lines into *Hh*-colonised Foxp3^GFP^ hosts for *in vivo* two-photon live imaging (Fig 2a). Host LA containing Tregs appear as green clusters. Transferred TCR^Hh^ are red, but co-express GFP upon TCR stimulation or IL-10 production. Acquisition in 3-dimensions allows for distinction of green/red co-expression versus two overlapping cells (Supp video 1).

**Figure 2.**
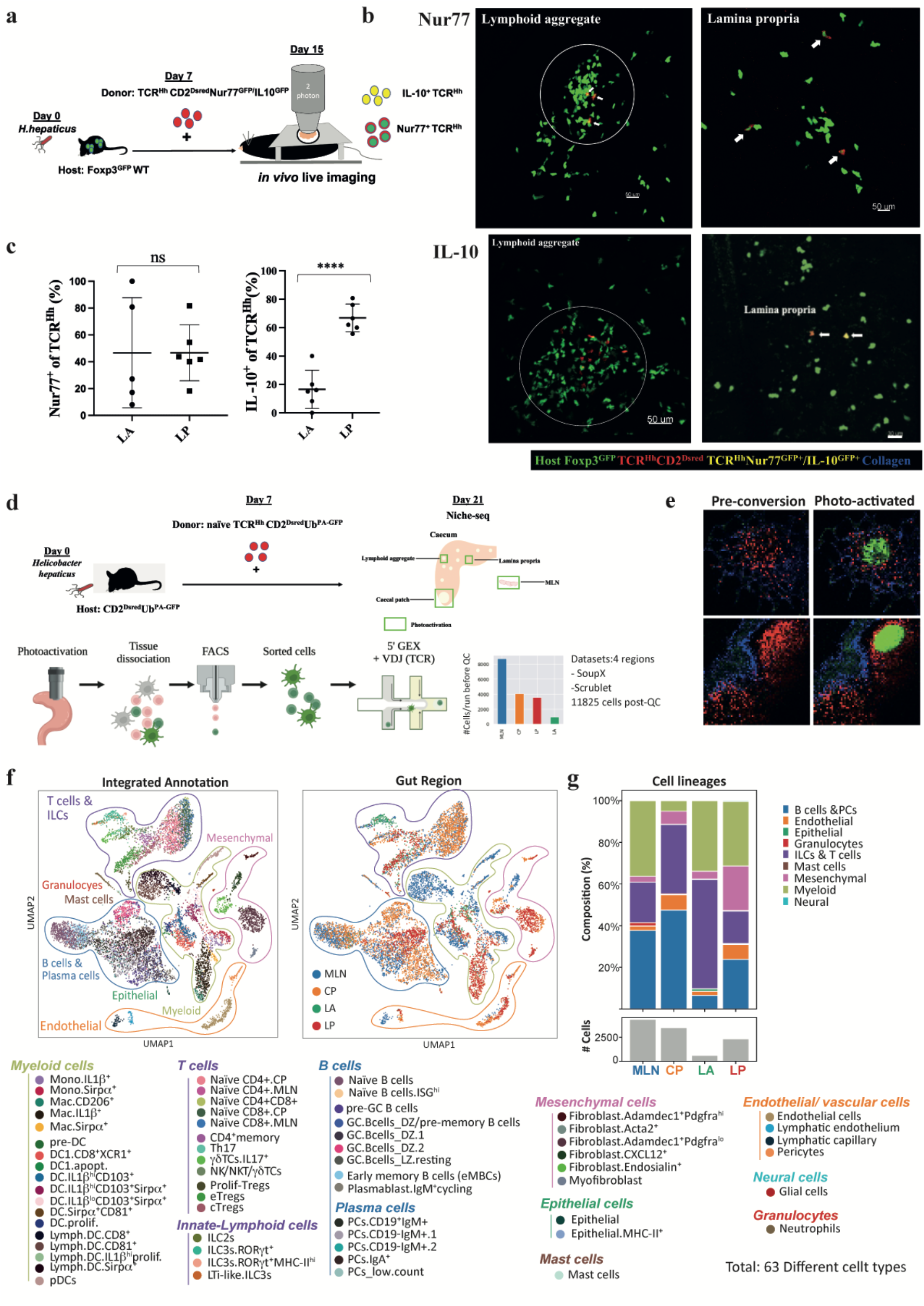
IL-10 production by TCR^Hh^ is enriched in the LP, and NICHE-seq allows identification of cellular composition across T cell microniches. a. Two photon *in vivo* live imaging of transferred naive TCR^Hh^CD2^dsRed^Nur77^GFP^ or TCR^Hh^CD2^dsRed^IL-10^GFP^ into *Hh*-colonised Foxp3^GFP^ hosts.
b. Representative images of TCR^Hh^Nur77^GFP^ cells (top panels) and TCR^Hh^IL-10^GFP^ (bottom panels) in the lymphoid aggregates (LA) and lamina propria (LP). TCR^Hh^ Nur77^GFP^/IL-10^GFP^ positive cells indicated by white arrowheads.
c. Frequency of Nur77^GFP^ (left) and IL-10^GFP^ (right) positive of TCR^Hh^ in the LA and LP.
d. NICHE-seq of gut tissue: Transfer of TCR^Hh^CD2^Dsred^Ub^PA-GFP^ into CD2^Dsred^Ub^PA-GFP^ hosts with subsequent photo-activation, cell sorting and scRNA-seq. Numbers of single cells acquired before and after QC as indicated.
e. Representative images of pre- and post-photoactivation of the LA (top panels) and T cell zone of CP(bottom panels) of CD2^Dsred^Ub^PA-GFP^ hosts.
f. UMAP visualisation of total cell subsets and detailed cell type annotations in each cell lineage (left) and their distribution in each gut location (right).
g. Bar chart showing cell subsets and total cell numbers at the indicated gut locations.

Within tissue, TCR^Hh^Nur77^GFP+^ T cells were present within LA (Fig 2b). Surprisingly, TCR^Hh^Nur77^GFP+^ cells were also located within the LP, spatially distant from LA (Fig 2b). This even distribution of recently activated T cells (Fig 2c) suggest TCR-MHCII interactions occur between TCR^Hh^ and LP-resident APCs, and/or rapid migration of TCR-stimulated TCR^Hh^ out of LA into LP after TCR engagement. Indeed, time-lapse videos reveal TCR^Hh^Nur77^GFP+^ cells to be highly motile (Supp video 2). We next wondered which tissue niche supported the highest production of IL-10 [12]. *In vivo* live imaging of donor TCR^Hh^CD2^Dsred^IL10^GFP^ in *Hh-*colonised Foxp3^GFP^ hosts demonstrated that IL-10 production by TCR^Hh^ cells was largely restricted to the LP (Fig 2b-c). IL-10^+^ TCR^Hh^ cells were actively motile throughout the tissue (Supp video 3), suggesting far-reaching suppressor function in response to *Hh*.

Because TCR^Hh^ cells that have recently experienced TCR stimulation are distributed throughout the LP and LA niches, whereas production of the Treg effector molecule IL-10 is highest in the LP niche, we wondered which cellular interactions and/or molecular cues in the LP niche drive TCR^Hh^ activation and Treg effector functions. We used two-photon photoactivation labelling of cells within T cell niches followed by scRNA-seq (NICHE-seq) to uncover transcriptional states and cellular composition of tissue microniches [1, 13]. Naïve T cells from TCR^Hh^ CD2^Dsred^ crossed to ubiquitously expressing photoactivatable GFP mice (TCR^Hh^ CD2^Dsred^Ub^PA-GFP^) [13] were transferred into *Hh*-colonised CD2^Dsred^Ub^PA-GFP^ hosts so both host and donor cells were photoactivatable. DsRed was used to visualise donor cells, and to mark T cell zones in host lymphoid tissue. We used two-photon microscopy to label donor and host cells with photoactivated GFP from four regions: the T cell zones of MLNs and CP, the LA and LP (Fig 2d-e and Supp Fig 5a). scRNAseq of GFP+ cells was performed using 10X and 5’ GEX and VDJ TCR sequencing.

A model built on previously published work [14–21] supported annotation of a spectrum of myeloid cells, T cells, ILCs, B cells, plasma cells, epithelial cells, mesenchymal cells, endothelial cells, and granulocytes across all 4 sites (Fig 2f-g, Supp Fig 5b, Supp Fig 6-7, and Supp Table 1). The relative distributions of these major intestinal cell types differ according to region, with a sizeable T cell population present in all micro-niches (Fig 2g). To understand how location affects suppressor function, we focussed on the lymphoid and myeloid populations in each niche that have the potential to form meaningful interactions with TCR^Hh^.

The T/ILC compartment comprises 16 transcriptionally distinct subsets (Fig 3a). As expected, naïve CD4^+^ and CD8^+^ T cells were mainly located in the MLNs and CP. ILC3s were dominant in the LA and ILC2s in the LP. The LP also contained a sizeable population of memory CD4 T cells. As suggested by the imaging data, Tregs were enriched in both LP and LA.

**Figure 3.**
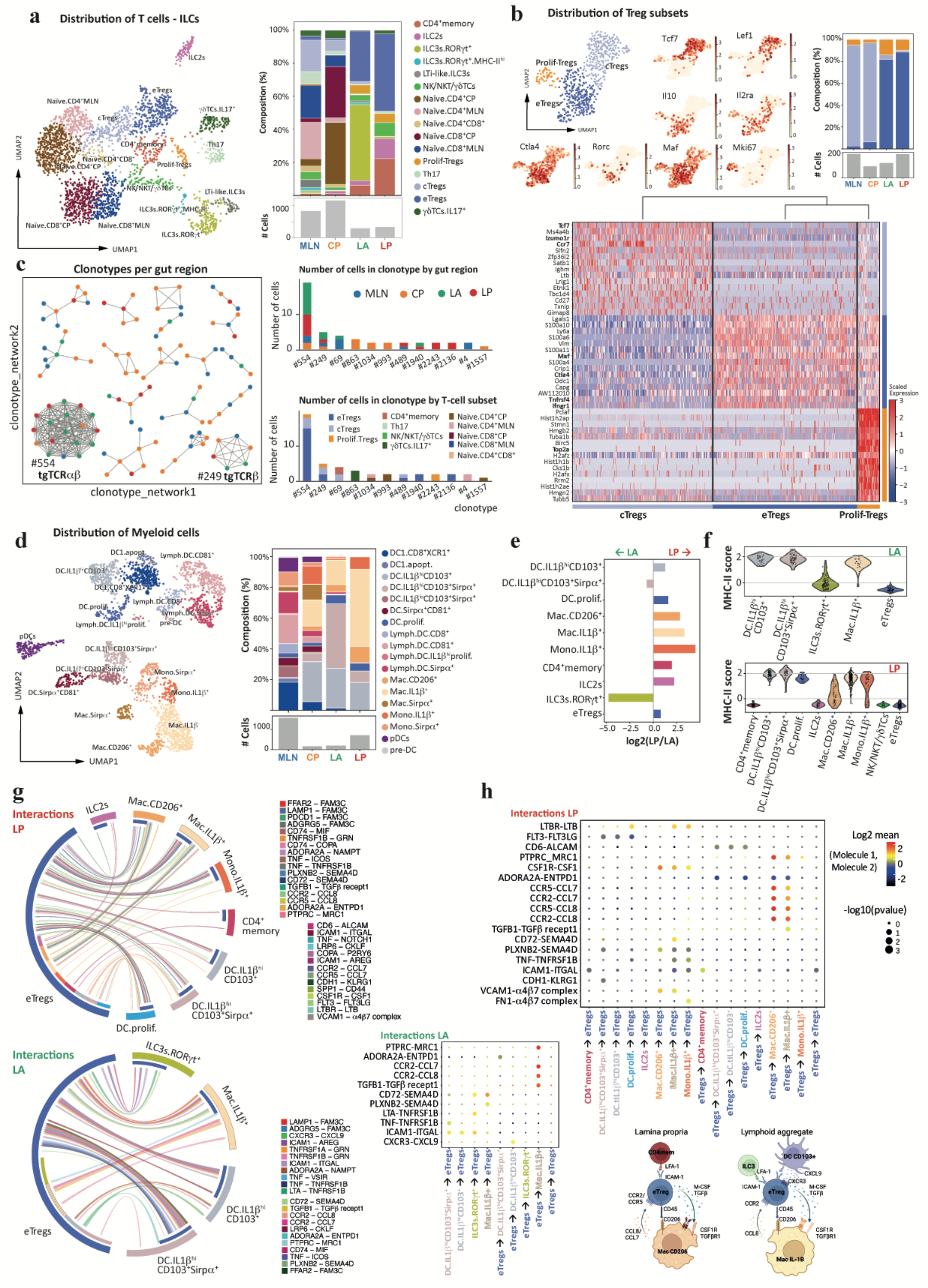
Lymphoid and myeloid cell subset data highlight distinct interactions in LP and LA microniches. a. UMAP visualisation of lymphoid subsets across all locations (left). Bar chart showing distribution of lymphoid subsets and cell numbers for each gut location (right). Total cell numbers at the indicated locations (bottom right).
b. UMAP visualisation of Treg subsets across all locations (top left) and overlay on the UMAP plot of the expression of selected genes (centre). Bar chart showing the distribution of Treg subsets and total cell numbers at the indicated locations (right). Heatmap of differentially expressed genes in the indicated Treg subsets across all locations (bottom).
c. Visualisation of clonotype network analysis of TCR clones (left). Barchart showing the distribution of TCR clones by gut location (right top) and T cell subset (right bottom).
d. UMAP visualisation of total myeloid subsets across all locations (left). Bar chart showing myeloid subsets and total cell numbers at the indicated locations (right).
e. Bar plot showing the fold change (log2) of proportions for myeloid, T cells and ILCs between LP and LA.
f. Violin plots of MHC-II expression score across lymphoid and myeloid cells in the LA and LP (restricted to cell types with >30 cells per region).
g. Chord diagrams showing significant interactions of Tregs with lymphoid and myeloid cells, separately for each gut location, LP (left) and LA (right), restricted to cell types with >30 cells per region.
h. CellPhoneDB analysis of receptor/ligand interactions in the LP (top) and LA (centre), restricted to cell types with >30 cells per region and excluding MHC-II interactions. Schematic summarising the most relevant cell-cell interactions (bottom).

Subclustering of Tregs revealed 3 subpopulations that are differentially distributed between SLOs and the tissue (Fig 2b). Lymphoid Tregs express *Tcf7* and *Ccr7* [22, 23], tissue Tregs express effector-associated molecules *Ctla4, Maf and Tnfrsf4*, and proliferating Tregs express histone genes (Fig 3b) [7, 24]. To highlight their distinct functional phenotype and spatial segregation, we refer to lymphoid associated Tregs as central Tregs (cTregs) and tissue resident Tregs as effector Tregs (eTregs).

To determine whether the host and transferred cells were comparable for downstream analysis, we performed a T cell clonotype analysis. TCR repertoire sequencing revealed the TCR^Hh^ clone (clonotype 554 and 249) as well as many host clonotypes to be present in all four regions and predominantly differentiate into the eTreg phenotype (Fig 3c). Host and TCR^Hh^ Tregs had similar mean expression of classical suppressor genes such as *Ctla4, Tigit*, and *Il10* (Supp Fig 8a), allowing combination of clonotypes for further analysis.

We next examined myeloid APC subset(s) across the tissue microniches, focusing especially on cells that may affect eTreg function in the tissue [25]. Mapping of monocyte/macrophage and dendritic cell subsets across the 4 regions did not reveal a unique APC population (Fig 3d); however, we identified an enrichment of IL-1β^hi^CD103^+^Sirpα^+^ dendritic cells within LA, whilst IL-1β^+^ and CD206^+^ macrophages and IL-1β+ monocytes were most abundant in the LP (Fig 3e).

By comparing APC populations present at >30 cells per microniche, we could see MHCII score for LP-resident IL-1β^+^ macrophages was comparable to IL-1β^hi^CD103^+^ and IL-1β^hi^CD103^+^Sirpα^+^ dendritic cells (Fig 3f, gene list: Supp Table 2). Taking abundance and MHCII expression into consideration, IL-1β^+^ macrophages dominate the T cell activation landscape in the LP, whilst IL-1β^hi^CD103^+^Sirpa^+^ dominate in the LA. MHCII^+^ ILC3s have been shown to regulate CD4^+^ T cell responses to gut commensals and maintain homeostasis in the MLN [26, 27]. LA are enriched for ILC3s, with a small subset expressing MHCII; however, this is a smaller fraction than present in the MLN (Fig 3f and Supp Fig 8b).

After defining the cellular composition, we sought to understand the potential cellular interactions at each location. We performed CellPhoneDB analysis [28] to understand the interactions between eTregs and T/ILC/myeloid cells for each tissue micro-niche based on populations with greater than 30 cells present. In the LP, eTregs are capable of interacting with all cell populations within the niche with many potential interactions across APC subsets (Fig 3g). This fits with their dynamic nature as observed previously with intravital microscopy (Supp video 3). LA interaction diagram demonstrates eTregs have more interactions with IL-1β+CD103+Sirp1α+ DCs, in particular (Fig 3g).

CellPhoneDB analysis of LA and LP catalogue key interactions between cells in each tissue microniche. Of particular interest are the chemokine/chemokine receptor pairs that bring APCs and eTregs together, TGFβ, and cell-cell contact molecules, which all differ between LP and LA (Fig 3h). Within the LP microniche, eTregs have the strongest *Ccr2/Ccl8* (Supp Fig 8c), *Ptprc/Mrc1*, and *Csf1/Csfr1* pathway interactions with CD206 macrophages suggesting these cells may play a dominant role beyond TCR engagement. Within the LA microniche, the *Ccr2/Ccl8, Ptprc/Mrc1*, and *CD72/Sema4d* instead connect eTregs with IL-1β+ macrophages, and *Cxcr3/Cxcl9, Icam1/Itgal* interactions connect eTregs with LA DC populations. In addition to differences in interactions with APCs, the eTregs in the LP had the strongest *Icam1/Itgal* interactions with CD4 memory T cells and *Ltbr/Ltb* interactions with ILC2s whereas eTregs in the LA could interact with ILC3s through *Icam1/Itgal* (Fig 3h).

Based on our imaging data (Fig 2b-c) and the differences in cell interactions within the tissue microniches, we suspected the LP and LA eTregs would have additional phenotypic differences. By comparing the eTreg populations across the tissue micro-niches, we found upregulation of *Gzmb, Areg, Ccr2*, and *Il2ra* in the LP microniche and *Cxcr4* in the LA (Fig 4a) [29, 30]. This highlights the LP as a key site that favours optimal eTreg function. This was also supported by mean expression data of known Treg effector molecules showing increases in *Areg, Gzmb, Icos, Tigit*, and *Tnfrsf18* (GITR) in the LP (Fig. 4a). Velocity pseudotime analysis of eTreg, cTreg, and proliferating Tregs suggests a trajectory of eTregs from MLN/CP to LA and on to LP with proliferating Tregs more closely related to eTregs (Fig 4b and Supp Fig8d-e). This raises questions about whether LP eTregs are terminally differentiated or whether these cells are plastic and capable of proliferating *in situ*.

**Figure 4.**
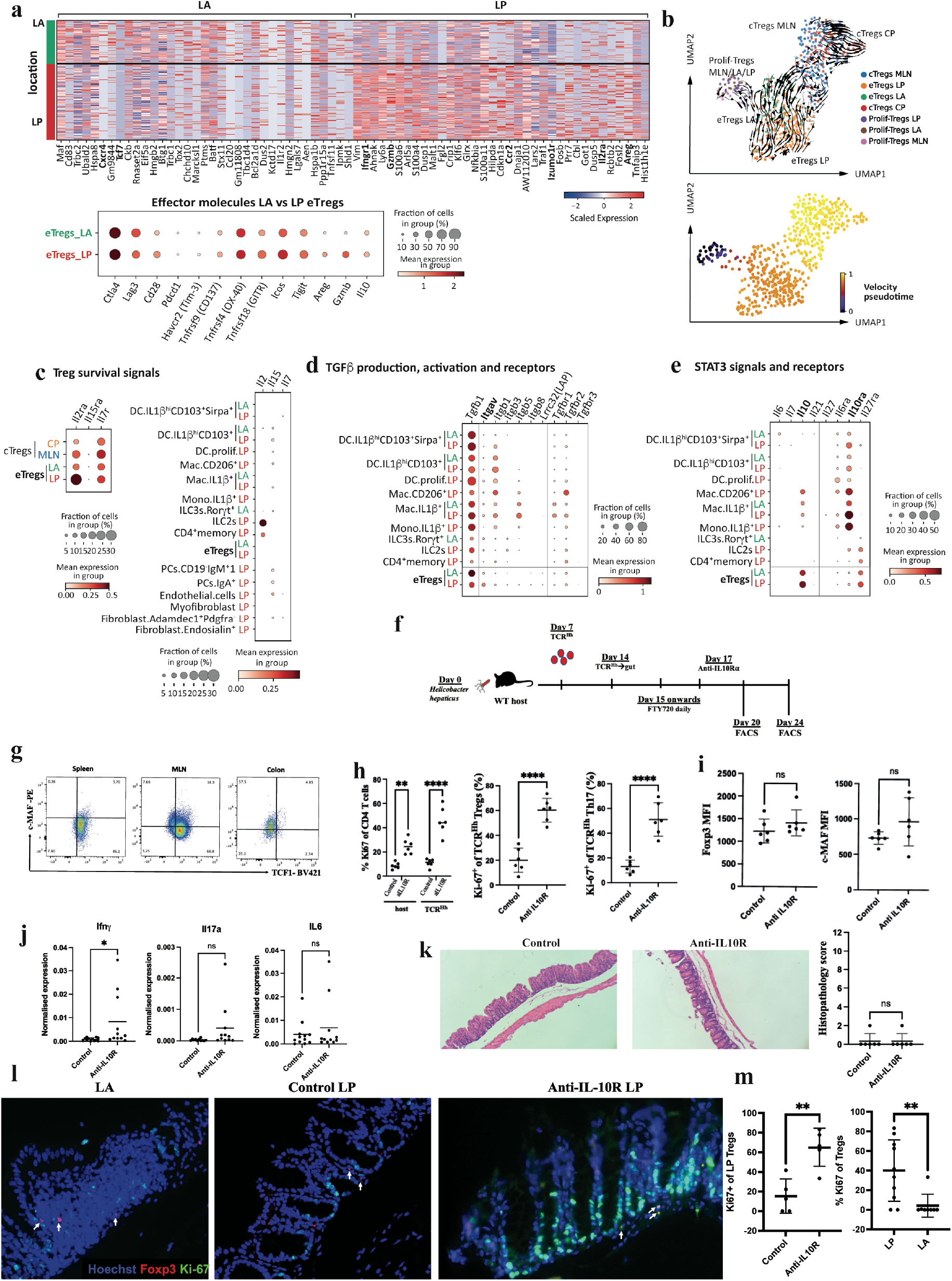
The LP niche promotes eTreg survival, phenotype, and function. a. Heatmap showing differential gene expression of eTregs in the LA *versus* LP (top) and dot plot showing relative expression of selected effector molecule genes within eTregs located in the LA *versus* LP (bottom).
b. UMAP of Treg subsets per location, filtering out the subsets with less than 10 cells in a given location (i.e. excluding eTregs_MLN, eTregs_CP, cTregs_LA, cTregs_LP and Prolif-Tregs_CP). Arrows depict summarised scVelo differentiation trajectories (above). Heat map overlay on the Treg UMAP indicating velocity pseudotime, as calculated by scVelo (below).
c. Dot plot showing relative expression of survival cytokine genes expressed by all the cell types with >30 cells in LA or LP regions (right) and their receptors in Treg populations (left).
d. Dot plot showing relative mean expression of TGF-β and its associated receptor and activating integrin genes in the main myeloid and lymphoid subsets (>30 cells per region) in the LA and LP.
e. Relative mean expression of STAT3 signalling genes and receptors within main myeloid and lymphoid subsets (>30 cells per region) in the LA and LP.
f. TCR^Hh^ were transferred into *Hh* colonised host mice, treated with daily FTY720 from day 15 and anti-IL10-Rα at day 17.
g. FACS validation of c-MAF and TCF1 protein expression in spleen, MLN and tissue sites of host Tregs at day 20 post *Hh* infection.
h. Ki-67 of total CD4 T cells of TCR^Hh^ and host cells (left), Ki-67 of TCR^Hh^ Tregs (centre) and Ki-67 of TCR^Hh^ Th17 cells (right) in control and anti-IL-10Rα treated mice at day 24.
i. MFI of Foxp3 (left) and c-MAF (right) of TCR^Hh^ Tregs from control and anti-IL10Rα treated mice at day 24.
j. Expression of *Ifng, Il17a* and *Il6* in caecum tissue by qPCR in control and anti-IL10Rα treated mice at day 24
k. Representative H and E staining (left) and histopathology score (right) of caecum tissue from control and anti-IL10Rα treated mice at day 24.
l. Representative immunofluorescent staining of Foxp3 and Ki-67 in LA (left panel, arrows mark Ki-67 negative Tregs), control LP (centre, arrows mark Ki-67 negative Tregs), and anti-IL10Rα treated LP (right panel, arrows mark Ki-67 positive Tregs).
m. Frequency of Ki-67 positive Tregs in the LP of control and anti-IL10Rα treated mice (left) and frequency of Ki-67 positive Tregs in the LP and LA in both control and anti-IL10Rα treated mice (right).

With all the data pointing to unique interactions and enhanced eTreg functions in the LP niche, we wanted to determine whether key known pathways such as survival cytokines and upregulators of Maf, TGFβ and STAT-3 cytokines, [14, 31, 32] are also spatially segregated within the tissue microniches. To understand which survival cytokines are required in the LP niche, we looked at receptors for gamma-chain cytokines on our Treg populations of interest. eTregs in the LP and LA express *Il2r* with highest expression in the LP microniche (Fig 4c). To determine which are the important cells that produce IL-2, we looked for expression in the cells present in the tissue. ILC2 and CD4 memory cells are the dominant sources of IL-2, with both populations present in the LP microniche and shown to have potential for interactions with eTregs in the LP (Fig 4c and Fig 3g).

TGFβ is widely expressed by most myeloid and lymphoid populations, including eTregs in the LA and LP (Fig 4d). However, eTregs in the LP are one of the few tissue populations that also express high levels of *Itgav* as well as *Itgb8* (Fig 4d), encoding the integrin subunits alpha V and beta 8 that are required to activate latent TGFβ [33]. This activated TGFβ can act in an autocrine and paracrine manner to control local immune responses, which is required for immune homeostasis [34, 35]. We next performed a comprehensive search of STAT3-dependent cytokine genes across the most abundant myeloid and lymphoid subsets in the tissue. We found upregulated gene expression of the pro-inflammatory cytokine *Il6* in IL-1β^+^CD103^+^Sirpα^+^ DCs resident in the LA (Fig 4e). In contrast, the LP was dominated by *Il10*,expressed primarily by eTregs with some expression in CD206^+^ and IL-1β^+^ macrophage subsets (Fig 4e). A small amount of *Il27* could be detected in the IL-1β^+^ macrophage population, specifically in the LP. Together these data establish myeloid cytokine microniches capable of tuning STAT-3 signals, potentially establishing inflammatory (LA) and anti-inflammatory (LP) niches within the intestinal tissue. We next examined expression levels of cytokine receptors. Moderate levels of *Il6ra* can be detected across myeloid populations, and *Il27ra* expression is largely restricted to eTreg, CD4+ memory T cells, and ILC2 populations. *Il10ra* is highly expressed on CD206^+^ and IL-1β^+^ macrophages and IL-1β^+^ monocytes, especially in the LP niche (Fig 4e). Macrophage sensing of IL-10 is critical for gut homeostasis [36, 37], and this data supports a positive feedback loop in the LP. CD206^+^ and IL-1β^+^macrophages respond to IL-10 and produce IL-10 and IL-27 [38, 39], activating STAT3 and supporting the *Maf* program in eTregs in the LP. This raises the question of whether acute perturbation of the IL-10 pathway could disrupt the LP eTreg phenotype and function.

After identifying spatially separated cTreg and eTreg responses to *Hh* and enhanced eTreg suppressor function within the LP, we sought to determine whether an inflammatory challenge that interferes with IL-10, a key eTreg effector molecule in the LP microniche, can affect eTreg phenotype or proliferation. We transferred TCR^Hh^ into colonised hosts as before. TCR^Hh^ migrate to the gut 7 days after transfer and adopt an IL10^+^ Treg fate (Fig 1d, f and h). To isolate the interactions in the gut tissue niches from the SLOs, FTY720 was injected daily from day 8, resulting in a pool of TCR^Hh^ cells trapped in the tissue (Fig 4f). We subsequently disrupted the IL10 positive feedback loop with an IL10Rα blocking antibody 10 days after cell transfer. Flow cytometry analysis validated NICHE-seq RNA expression of *Tcf7* in cTregs and *Maf* expression in eTregs (Fig 4g). At 3 days after anti-IL10Rα treatment (13 days after TCR^Hh^ cell transfer), there was no significant difference between the two treatment groups in terms of TCR^Hh^ Treg proportion or phenotype (Supp Fig 9a). However, by 7 days after anti-IL10Rα treatment (17 days after TCR^Hh^ cell transfer), there was an increase in local proliferation of TCR^Hh^ and host T cells with approximately 60% of TCR^Hh^ Tregs and Th17s expressing Ki-67 (Fig 4h). It has been suggested that eTregs may be terminally differentiated and therefore less able to proliferate; however, proliferating eTregs expressed similar levels of Foxp3 and C-maf compared to controls (Fig 4i and Supp Fig 9b). 7 days after anti-IL10Rα treatment, while we observe some increased cytokine expression, we did not observe overt inflammation (Fig 4j-k and Supp Fig 9c) suggesting that while IL-10 is critical to restrain Th17s, other eTreg suppressor functions are able to constrain the local inflammatory response. To determine which niche in the intestinal tissue supported Treg proliferation in this perturbed environment, we stained tissues with Ki67, Foxp3, and Hoechst. This demonstrated Treg proliferation in the LP niche (Fig 4l-m), again supporting the LP as the key location for eTreg function in the gut. These findings taken together with our Niche-seq data suggest that eTreg suppressor functions (e.g. Tigit, ICOS, and GITR Fig 4a and Supp Fig 10) in the LP can constrain the inflammatory response when the local IL-10 circuit is disrupted.

## Discussion

In this study, we follow the natural history of microbe-reactive Tregs as they acquire and maintain immune regulatory function in the intestine. By examining cells in anatomical microniches, we have revealed spatial diversity of cTreg and eTreg subsets in lymphoid and non-lymphoid tissues and the importance of the lamina propria niche, not the lymphoid aggregates, as the site of enhanced eTreg function. The LA microniche may act as an entry point for eTregs into the tissue with the potential to skew the response toward inflammation through eTreg interactions with DCs and ILC3s. The LP niche is unique in its myeloid cell makeup with interactions between macrophages and eTregs through antigen presentation, CCR2/CCL8, CD45/CD206, and IL10/IL10R axes contributing to recruitment of highly motile eTregs and fine tuning of the immune response to pathobiont colonization.

By perturbing the system with IL-10Ra blocking antibodies, we demonstrate that eTregs are able to maintain their c-Maf and Foxp3 levels while proliferating in LP niche in the context of an inflammatory milieu. While IL-10Rα blockade results in proliferation of Th17 cells, colitis does not develop, likely due to additional eTreg effector molecules including GITR, Tigit, CTLA-4, and granzymeB that remain intact.

Cell therapy and vaccination strategies to promote and establish tolerance in inflammatory diseases will rely on Tregs that can function in inflammatory conditions, and we provide support for eTregs as an ideal therapeutic population, capable of maintaining their phenotype in an inflammatory milieu. Information gained from the study of microniches can be used for targeted interventions to support eTreg recruitment, activation, differentiation, survival and function in the inflammatory niche within the intestine and beyond.

## Author contributions

Yisu Gu designed, performed, and analysed the experiments and prepared the manuscript. Raquel Bartolomé-Casado conceived of and performed the single-cell sequencing analysis and prepared the manuscript. Chuan Xu created the analysis model and assisted with manuscript preparation. Alina Janney performed experiments. Cornelia Heuberger performed experiments. Claire Pearson facilitated and supervised experimentation and provided intellectual input. Sarah Teichmann supervised analysis and manuscript preparation. Emily Thornton designed, performed, and analysed experiments, supervised the project, and prepared the manuscript. Fiona Powrie designed experiments, and supervised the project and manuscript preparation.

## Acknowledgements

We would like to thank the Kennedy Institute of Rheumatology (KIR) Flow Cytometry Facility and the manager, Jonathan Webber, for help with flow cytometry and FACS, the KIR Biomedical Services Unit, especially Luke Barker for help with animal care and husbandry, and the KIR microscopy facility and manager Christoffer Lagerholm. We acknowledge the generous support of the Kennedy Trust for Rheumatology Research, IDRM and Carl Zeiss GMBH for the microscopy facilities (Zeiss 980) used in this research. We acknowledge the generous support of the Kennedy Trust for Rheumatology Research and a Wellcome Trust Multi-User Equipment Grant 202911/Z/16/Z for the microscope purchase (Zeiss 880 multiphoton) and facilities used in this research. We want to thank Dr Ida Parisi, Dr BryonysStott and Miss Rhiannon Cook in the Kennedy Institute of Rheumatology Histology Service for tissue processing and staining. We thank Angela Lee and Moustafa Attar in the Oxford Genomics Centre at the Wellcome Centre for Human Genetics (funded by Wellcome Trust grant reference 203141/Z/16/Z) for the generation and initial processing of sequencing data. We also acknowledge the support received from Stijn van Dongen at the Wellcome Sanger Cellular Genetics Informatics team and the Teichmann lab for discussion and support with scripts.

YG was funded by a Wellcome Trust Clinical Research Fellowship (CRTF), grant reference 201224/Z/16/Z.

RB-C Grant 315307, Forskerprosjekt 2020, Researcher Project / International Mobility Grant from the Research Council of Norway and travel grant from the Per Brandtzæg’s Fund for Research in Mucosal Immunology.

ET was supported by Wellcome Trust (095688/Z/11/Z and 212240/Z/18/Z, awarded to FP), Nuffield Department of Medicine, and MRC core grant reference MC_UU_00008.

FP was supported by Wellcome Trust (095688/Z/11/Z and 212240/Z/18/Z).

## Competing interest statement

FP receives consultancy fees or research support from Janssen, GSK, T-Cypher and Novartis

In the past three years, S.A.T. has consulted or been a member of scientific advisory boards at Roche, Genentech, Biogen, GlaxoSmithKline, Qiagen and ForeSite Labs and is an equity holder of Transition Bio.

## Notes

### Summary of Updates

Updated to correct a typo in the affiliations.

